# Purinergic Signaling Drives Multiple Aspects of Rotavirus Pathophysiology

**DOI:** 10.1101/2024.05.07.592953

**Authors:** Kristen A. Engevik, Francesca J. Scribano, J. Thomas Gebert, Joseph M. Hyser

**Author notes:** **Disclosures:** The authors declare that the research was conducted in the absence of any commercial or financial relationships that could be construed as a potential conflict of interest. **Data Availability:** The data that support the findings of this study are available from the corresponding author, JMH, upon reasonable request. **Author Contributions**: Concept and design (KAE, JMH); intellectual contribution (KAE, FJS, JTG, JMH); data acquisition (KAE, FJS, JTG); data analysis, statistical analysis, and interpretation (KAE); drafting manuscript (KAE); obtained funding (KAE, FJS, JTG, JMH). **Corresponding Author:** Joseph Hyser, PhD. 1 Baylor Plaza Houston TX 70030.

## Abstract

Rotavirus causes life-threatening diarrhea in children, resulting in ∼200,000 deaths/year. The current treatment during infection is Oral Rehydration Solution which successfully replenishes fluids but does not alleviate diarrhea volume or severity. As a result, there is an urgent need to better understand rotavirus pathophysiology and develop more effective pediatric therapeutics. Rotavirus primarily infects the tips of small intestinal villi, yet has far-reaching effects on cell types distant from infected cells. We recently identified that rotavirus infected cells release the purinergic signaling molecule ADP, which activates P2Y1 receptors on nearby uninfected cells *in vitro*. To elucidate the role of purinergic signaling via P2Y1 receptors during rotavirus infection *in vivo*, we used the mouse-like rotavirus strain D6/2 which generates a severe infection in mice. C57BL/6J mouse pups were given an oral gavage of D6/2 rotavirus and assessed over the course of 5-7 days. Beginning at day 1 post infection, infected pups were treated daily by oral gavage with saline or 4 mg/kg MRS2500, a selective P2Y1 antagonist. Mice were monitored for diarrhea severity, diarrhea incidence, and viral shedding. Neonatal mice were euthanized at days 3 and 5 post-infection and small intestine was collected to observe infection. MRS2500 treatment decreased the severity, prevalence, and incidence of rotavirus diarrhea. Viral stool shedding, assessed by qPCR for rotavirus gene levels, revealed that MRS2500 treated pups had significantly lower viral shedding starting at day 4 post infection compared to saline treated pups, which suggests P2Y1 signaling may enhance rotavirus replication. Finally, we found that inhibition of P2Y1 with MRS2500 limited transmitted rotavirus diarrhea to uninfected pups within a litter. Together, these results suggest that P2Y1 signaling is involved in the pathogenesis of a homologous murine rotavirus strain, making P2Y1 receptors a promising anti-diarrheal, anti-viral therapeutic target to reduce rotavirus disease burden.

## Introduction

Diarrheal diseases are a major cause of morbidity in children under 5 years old with approximately 446,000 deaths globally each year^1^. The majority of cases are due to enteric viruses, which cause diarrhea in fundamentally different ways than bacteria^1–3^. Rotavirus is the leading cause of severe gastroenteritis globally in infants and children^4^, with approximately 258 million episodes of diarrhea in children and 200,000 deaths annually^1, 5^. The increase in rotavirus vaccination since 2006 has significantly reduced death and incidence in high income countries^5–8^, however live attenuated vaccines in low and middle income countries have been less effective in reducing severe rotavirus disease burden and mortality^5–7, 9^. Rotavirus diarrhea and vomiting cause potentially fatal dehydration, and can impair long-term childhood development^10^. Furthermore, as with other pediatric diarrheal diseases, there are currently no approved anti-secretory drugs nor anti-motility drugs (e.g., Imodium) suitable for children due to the risk of ileus^10, 11^. Thus, elucidating the signaling pathway(s) that rotavirus exploits to induce diarrhea represents an opportunity to better inform and develop treatments for rotavirus infection in children.

Rotavirus infects enterocytes located at the tips of small intestinal villi, yet rotavirus infection is associated with wide-raging effects throughout the intestine that involve other cell types^12–14^. During infection, rotavirus enters enterocytes and expresses both nonstructural proteins that aid in replication and structural proteins that assemble into progeny virions. Nonstructural protein 4 (NSP4) is one of the rotavirus nonstructural proteins that has been shown to be a key regulator of virus replication and a virulence factor involved in rotavirus diarrhea. Within virus-infected cells, intracellular NSP4 functions as a viral ion channel to release Ca^2+^ from the endoplasmic reticulum, which initiates the reprogramming of cells for virus replication. Further, the NSP4-induced increase in Ca^2+^ signaling in infected cells also induces the release of paracrine signals, which affect neighboring uninfected cells^15–20^. We recently showed that rotavirus infected cells exhibit robust Ca^2+^ signaling, which drives increased extracellular purines that activate the purinergic receptor P2Y1 on neighboring cells, and results in a propagating “intercellular Ca^2+^ wave” encompassing neighboring cells^21^. We propose that these intercellular Ca^2+^ waves drive several aspects of the cell-cell communication observed in rotavirus infection, allowing for only a few infected cells to amplify the functional dysregulation of the gastrointestinal tract. We hypothesized that blocking P2Y1 *in vivo* would attenuate hallmarks of infection, including diarrhea and disruption of the intestinal architecture. Herein we demonstrate that P2Y1 signaling drives several aspects of rotavirus infection, including duration of infection, severity of rotavirus driven-diarrhea and rotavirus shedding.

## Materials & Methods

### Mouse Model of Rotavirus Infection

All animal experiments were performed with approval by the Institutional Animal Care and Use Committee (IACUC) at Baylor College of Medicine, Houston, TX, in accordance U.S. National Institutes of Health guidelines. For all mouse experiments, we used natural litters with both female and male animals, as there are no known sex-mediated effects of rotavirus infection or diarrhea^22^. C57BL/6 mice were purchased from Charles River Laboratory, maintained on a normal mouse diet (7922 NIH-07 Mouse diet, Harlan Laboratories, Indianapolis, IN) and bred in a BSL2 facility. Dams and natural litters were randomly allocated into groups (n=6-11) at day 4-7 of life. The groups were uninfected, rotavirus infected with saline treatment, or rotavirus infected with MRS2500. Murine-like (D6/2) rotavirus was acquired from Dr. Harry Greenberg from Stanford University. Pups were orally gavaged with either 50 µl of 2.5×10^5^ FFU murine-like D6/2 rotavirus (a dose 25x the DD50), which is a reassortant between murine EW and similar RRV rotavirus strains^23^, or the same volume of MA104 lysate. Inoculum of D6/2 rotavirus or MA104 lysate was mixed with sterile-filtered green food dye (3% by volume) for visualization of the gavage. The following day, mice were provided daily oral gavage with 4 mg/kg MRS2500 or the equivalent volume of vehicle control (sterile saline).

For transmission studies, CD-1 dams with natural litters (purchased from Center for Comparative Medicine, Baylor College of Medicine) were used and housed at a BSL2 facility in standard cages with normal mouse diet and water provided ad libitum. Dams and natural litters were randomly allocated into groups (n=9-14) at day 4 of life. In each cage, 30% of littermates were inoculated with 50 µl of 1×10^4^ FFU murine-like (D6/2) rotavirus^23^ by oral gavage, while the remaining 70% of littermates were received oral gavage of 4 mg/kg MRS2500 or the equivalent volume of vehicle control (sterile saline) on the same day. MRS2500 or sterile saline were given daily to uninfected littermates from 0-6 days post infection.

Each day post infection, pups’ abdomens were gently palpated to express feces. Stools were graded on a 4-point scale for volume, consistency, and color as described previously^24, 25^. Diarrhea is defined as an average score of ≥2. Diarrhea incidence was calculated as the number of stools with a diarrhea score ≥ 2 divided by the number of stools collected per cage per day. Overall diarrhea prevalence was calculated as the percentage of pups within a litter that experienced diarrhea at any point during the experiment. Pups from which a stool sample could not be collected were not counted in the diarrhea severity or prevalence calculations. Animals were euthanized on day 1, 3, and 5 post-D6/2 rotavirus infection.

### Tissue Fixing and Staining

Excised intestinal segments were fixed in 10% formalin (Sigma Aldrich, #HT501128) overnight. The tissue was collected from RV infected pups treated with PBS or MRS2500 irrespective of diarrhea scores. The tissue was embedded in paraffin and 7-µm sections were used for Hemotoxylin & Eosin (H&E) and immunostaining. Briefly, H&E stains were performed by a series of incubations with alum haematoxylin, 0.3% acid alcohol, Scott’s tap water and eosin and slides were dehydrated, and a coverslip applied with mounting media (ThermoFisher, #14-390-5). For immunostaining, slides were dehydrated, and antigens were exposed by incubating slides in Vector Labs Antigen Unmasking Solution Citrate Buffer pH 6 (Vector labs, #F4680) in a steamer for 30 min. Sections were blocked for 1 h at room temperature in 10% donkey serum. Sections were incubated with primary antibodies (**Table 1**) overnight at 4°C, followed by secondary antibodies (**Table 1**) for 1 hr at room temperature and incubation with Hoechst 33342 (Sigma, #B2261) at room temperature for 10 min. All sections were cover-slipped with Fluoromount Aqueous Mounting Medium (Sigma Aldrich, #F4680). Slides were imaged with widefield epifluorescence on a Nikon TiE inverted microscope using a SPECTRA X LED light source (Lumencor) using a 20X Plan Apo (NA 0.75) objective. Color images for H&E sections were captured using a DS-Fi1-U2 camera (Nikon). Fluorescence images were captured using an ORCA-Flash 4.0 sCMOS camera (Hamamatsu). Quantitative analysis of fluorescent stains was accomplished using FIJI (Formerly ImageJ) software by tabulating mean pixel intensity (National Institutes of Health) as previously described ^26^ or manually counting number of fluorescent stain cells.

**Table 1:** Primary and Secondary Antibodies.

| Type | Target | Species | Recognizes | Company | Category | Dilution |
| --- | --- | --- | --- | --- | --- | --- |
| Primary | E-cadherin | Rabbit | Mouse, human | GeneTex | GTX100443 | 1:100 |
| Primary | Rotavirus | Guinea Pig | Mouse, human | In-house | In house antibody | 1:1000 |
| Secondary | Anti-rabbit Alexa Fluor 488 | Donkey | Rabbit | Rockland | 611-741-127 | 1:500 |
| Secondary | Anti-rabbit Alexa Fluor 555 | Donkey | Guinea Pig | Rockland | 606-142-129 | 1:500 |

### qPCR

RNA from feces was extracted according to manufacturer details (Invitrogen, # 15596026). The SensiFAST cDNA synthesis kit (Bioline USA Inc., # BIO-65053) was used to synthesize cDNA from 1 µg RNA. Fecal rotavirus levels were examined by quantitative real-time PCR (qPCR) using Fast SYBR Green (Thermo-Fisher, #4385617) and primers designed with PrimerDesign Software on a Qunatstudio3 qPCR Machine (ThermoFisher) (**Table 2**). Relative fold change was calculated using the ΔΔCT with the housekeeping gene 18S.

**Table 2:**
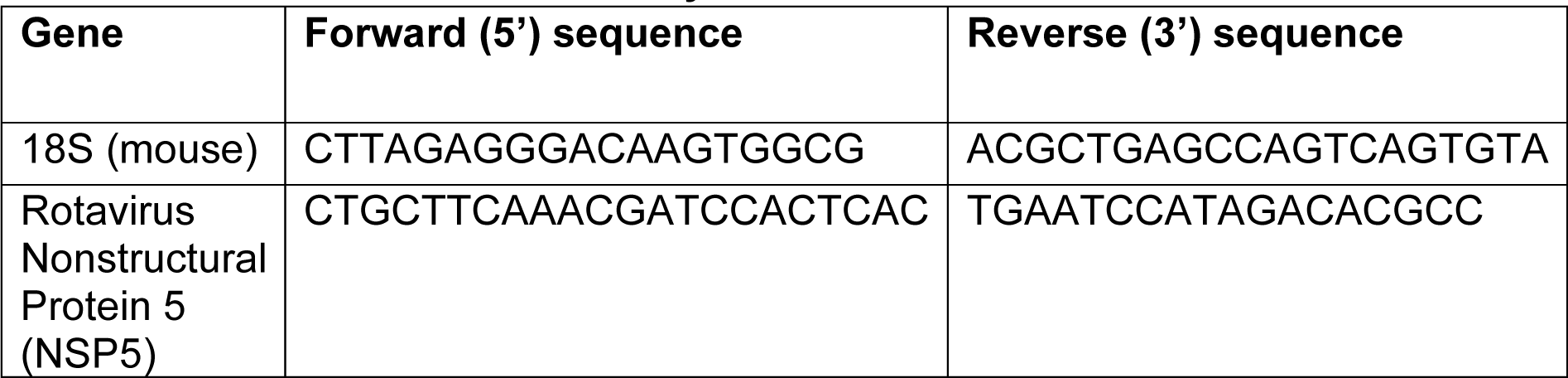
Primers Used in This Study.

### Fluorescence Foci Assay

To assess shedding of rotavirus in stool collected from CD-1 pup transmission studies, we utilized a fluorescence foci assay similar to previously described protocols^21, 27, 28^. In brief, stool samples were homogenized in 40µL of PBS with calcium and magnesium. Then 10µL of homogenized stool samples were resuspended into 990µL DMEM media and supplemented with 10 µg/mL Worthington’s trypsin, followed by an incubation for 30-45 min at 37°C. Following incubation, stool samples were serially diluted 2-fold and incubated with MA104 cells (2.5×10^4^ cells/well in a 96-well plate) for 1 hour at 37°C. Following incubation, the inoculum was replaced with DMEM and incubated for 18-20 hours at 37°C. Monolayers were fixed with cold methanol for 20 minutes at 4°C, followed by three washes with 1x PBS and incubation with guinea-pig primary antibodies against RV (1:10,000) overnight at 4°C. Cells were then washed with 1X PBS three times and incubated with secondary antibody donkey anti-guinea pig AlexaFluor 549 (1:1,000) for 1 hour at 37°C kept in the dark, followed by three washes with 1X PBS before imaging. Quantitative analysis of fluorescent stains was accomplished using FIJI software by counting number of fluorescent cells.

### Cell Lines, Mouse Intestinal Organoid Cultures and Viruses

African green monkey kidney epithelial (MA104) cells were cultured in high glucose DMEM supplemented with 10% fetal bovine serum (FBS) at 37°C in 5% CO_2_. MA104 cells were engineered to stably express the GCaMP6s cytosolic calcium indicator (MA104-GCaMP6s) as previously described and maintained under 50 µg/mL hygromycin drug selection^20, 21^. D6/2-rotavirus was propagated in MA104 cells in serum-free DMEM supplemented with 1 µg/mL Worthington’s Trypsin (Worthington Biochemical). Harvested cells were then subjected to three freeze/thaw cycles. Mock or rotavirus-infected MA104 cell lysates were treated with 10 µg/mL Worthington’s trypsin, incubated for 30 min at 37°C, then used to inoculate MA104-GCaMP6s cells or mouse organoids.

For live imaging, MA104-GCaMP6s cell monolayers were generated by seeding 5 x 10^4^ cells into 8-well ibidi treated chamber slides (ibidi) and grown for 2-3 days until confluent. Mouse jejunum intestinal organoids were generated from BALB/c mice as previously described^26^. Organoids were grown in complete media with growth factors and maintained in culture in Matrigel (Corning, #354248). To infect organoids, lysate from D6/2-rotavirus infected MA104 cells was added to the organoid containing wells. As a control, uninfected MA104 lysate was added to organoid containing wells. The inoculum was removed after 1 hr and replaced with fresh media.

#### Microscopy and image analysis

For live calcium imaging of MA104-GCaMP6s cells, monolayers were placed in Okolabs stage-top incubation chamber with CO_2_ and humidity control. The stage was then placed on a Nikon TiE inverted widefield epifluorescence microscope (Nikon) with a motorized X, Y, and Z stage for software-controlled multi-position imaging. FluoroBrite DMEM supplemented with 15 mM HEPES, 1X sodium pyruvate, 1X GlutaMAX, and 1X non-essential amino acids was used for fluorescence calcium imaging. Cell images were obtained by widefield epifluorescence using a 20x Plan Apo (NA 0.75) objective, using a SPECTRA X LED light source (Lumencor). Calcium waves were quantified using a previously reported method in ImageJ^29^.

For organoid live imaging, mouse jejunum intestinal organoids were placed in Okolabs stage-top incubation chamber with CO_2_ and humidity control. The stage was then placed on a Nikon TiE inverted widefield epifluorescence microscope (Nikon) with a motorized X, Y, and Z stage for software-controlled multi-position imaging. Organoids were incubated overnight at 37°C with 5% CO_2_ and swelling was assessed by live imaging on a Nikon TiE microscope. Organoids were imaged using a SPECTRAX LED light source (Lumencor) and a 10x Plan Fluor phase contrast objective. Swelling was quantified in ImageJ as previously described^21^.

## Results

We have previously shown *in vitro* that human and simian rotavirus induce intercellular calcium waves through purinergic P2Y1 signaling over the course of infection in MA104 monkey kidney epithelial cells (MA104) and human intestinal organoid lines stably expressing the cytosolic calcium indicator, GCaMP6s^21^. As murine rotaviruses, and their NSP4 genes, are phylogenetically distinct from human and simian rotaviruses^30^, we tested whether the murine-like rotavirus D6/2 reassortant was capable of eliciting intracellular calcium waves in MA104-GCaMP6s cells. D6/2 rotavirus infected cells exhibited significantly more intercellular calcium waves than mock infected cells over the course of the infection (∼18-20 hours) (**Figure 1A, B**). Furthermore, we observed that addition of apyrase (an ATP-diphosphohydrolase) or BPTU (a P2Y1 receptor inhibitor) following D6/2 rotavirus infection significantly decreased the number of rotavirus-induced intercellular calcium waves (**Figure 1A, B**). These data indicate that, like human and simian rotaviruses, the murine-like D6/2 rotavirus strain induced paracrine intercellular calcium waves mediated by P2Y1 purinergic signaling, and further suggests that rotavirus’ ability to elicit intercellular calcium waves is conserved across Group A rotavirus strains. With confirmation that murine rotaviruses also generate intercellular calcium waves, we next investigated the role of P2Y1 signaling in a neonatal mouse model of rotavirus infection and disease.

**Figure 1:**
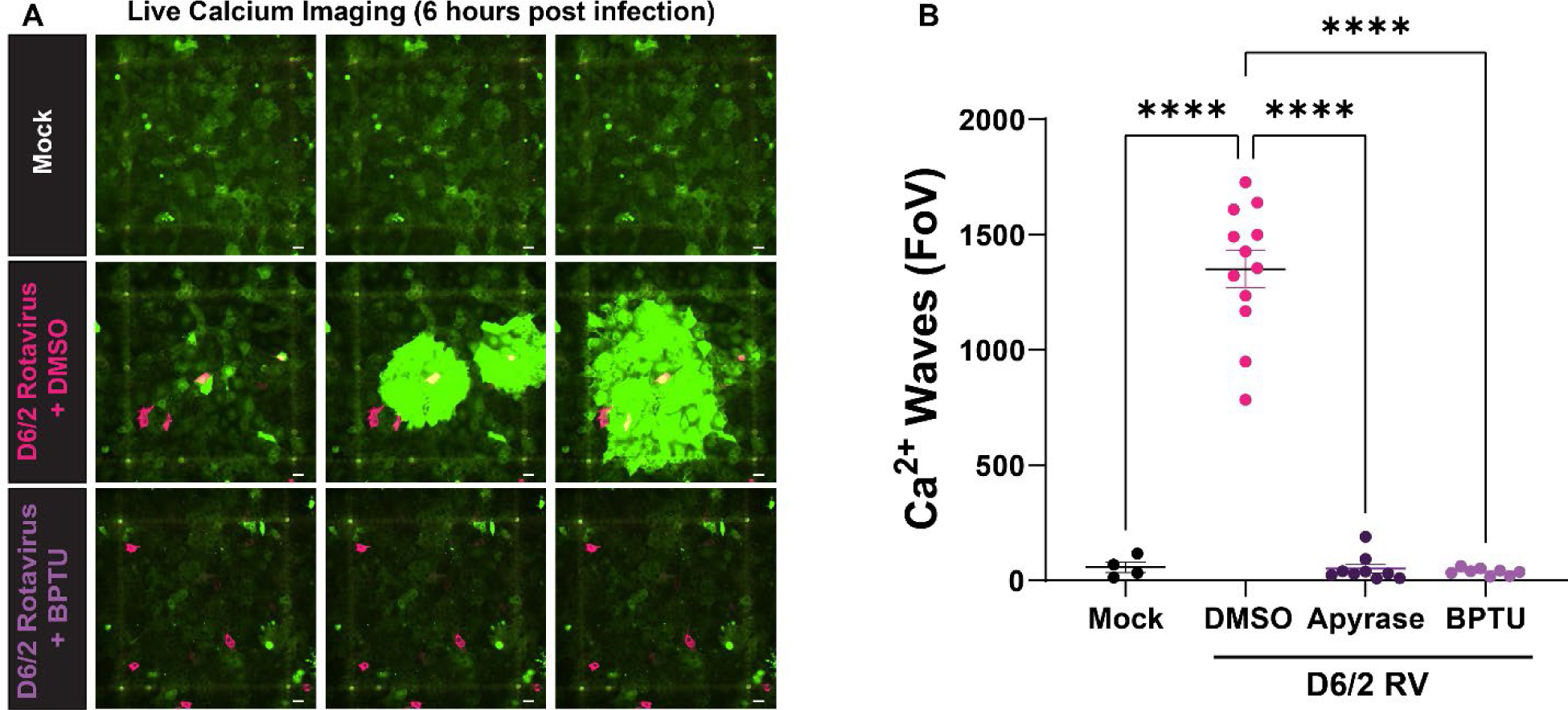
Mouse-like D6/2 rotavirus elicits intercellular calcium waves via purinergic P2Y1 signaling. **A**. Representative images of D6/2 RV infection of MA104-GCaMP6s cells (green), which exhibit intercellular calcium waves which propagate from infected cells (magenta) to neighboring uninfected cells. **B.** Quantification of the number of intercellular calcium waves during D6/2 RV infection in the absence or presence of apyrase (hydrolyzes extracellular ADP and ATP) or BPTU (P2Y1 inhibitor). One-Way ANOVA. ****p<0.0001. Scale bar = 50 µm.

Infant mice exhibit an age-dependent susceptibility to rotavirus diarrhea (≤14 days of age), which make them a model system that mimics human infant infection^31^. For these studies, neonatal C57B6 mice were orally gavaged with D6/2 rotavirus stock (2.5×10^5^ FFU) or an equal volume uninfected MA104 cell lysate as the negative control. In this model, diarrhea peaks 3-4 days post infection (DPI) and returns to baseline by 7 DPI. To examine the effects of P2Y1 inhibition on rotavirus diarrhea, from 1-6 DPI, mice received daily gavage of saline (vehicle control) or 4 mg/kg MRS2500, a selective P2Y1 competitive antagonist (**Figure 2A**) and diarrhea severity and incidence were monitored daily. Uninfected control mice never exhibited diarrhea with stool scores never exceeding an average of 1 (**Figure 2B, C, D**). In the rotavirus-infected groups at 2 DPI, 88.3% of the saline-treated pups exhibited diarrhea, but this was reduced to 61.8% of MRS2500 treated pups with diarrhea (**Figure 2B**). A lower incidence of diarrhea in the MRS2500-treated pups continued through 5 DPI (**Figure 2B**). In terms of severity, the rotavirus infected pups with PBS treatment exhibited a peak average diarrhea score of 2.2 ± 0.3 on 3 DPI compared to 1.8 ± 0.4 for the MRS2500-treated pups (**Figure 2C**). Similar to the incidence data, the trend of lower diarrhea scores continued with MRS2500 treated pups on 4 DPI and 5 DPI. Over the course of infection, the diarrhea prevalence, calculated by the number of pups within each litter that exhibited diarrhea at any point, was 95% diarrhea for PBS-treated pups, which was significantly higher than the 67% diarrhea prevalence observed in pups treated with MRS2500 (**Figure 2D**). Overall, blocking P2Y1 signaling with MRS2500 significantly decreased rotavirus diarrhea prevalence and severity, and resulted in faster resolution of disease by ∼2 days.

**Figure 2.**
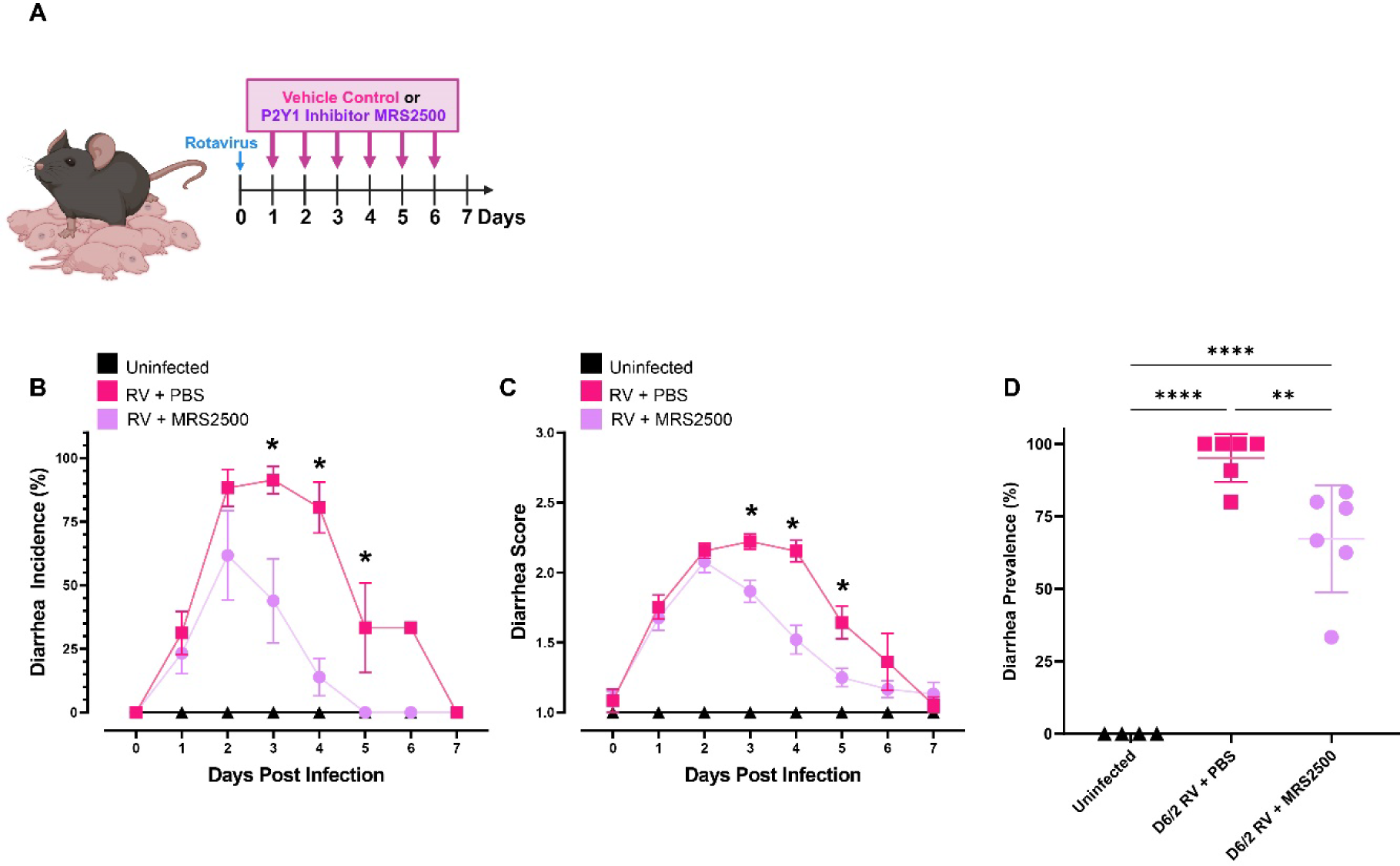
P2Y1 inhibition limits rotavirus diarrhea in vivo. **A**. Schematic of D6/2 rotavirus infection of C57B6/J pups. In C57B6/J dams with natural litters, pups were orally gavaged with D6/2 rotavirus at day 5-7 of life and monitored daily for diarrhea severity. Starting at 1 day post infection, litters were given either vehicle control (PBS) or 4mg/kg MRS2500 (P2Y1 inhibitor) by oral gavage daily. Stool and diarrhea scores were collected and assessed daily prior to treatment. **B**. The average percentage of diarrhea incidence each day in C57B6/J pups mock infected (black, n=4 litters) or D6/2 rotavirus infected, treated with PBS (n=6 litters, magenta) or MRS2500 (n=6 litters, purple). One-Way ANOVA, *p <0.05. **C**. Diarrhea scores over a 7-day period in mock infected pups (black) and D6/2 rotavirus infected pups treated with PBS (magenta) or MRS2500 (purple). One-Way Repeated Measures ANOVA, *p <0.05. **D**. Diarrhea prevalence over 7-day period in uninfected (black) and rotavirus infected pups treated with PBS (magenta) or MRS2500 (purple). One-Way ANOVA. **p <0.005, ****p <0.0001. n= 4-6 litters/group.

Rotavirus infection is known to stimulate chloride (Cl^-^) secretion, which contributes to the diarrhea observed in mice^32, 33,34, 35^. It has previously been shown in 3D human intestinal organoids that rotavirus infection results in significant increased swelling, due to virus-induced stimulation of fluid secretion (ie Cl^-^ secretion)^21^. Since our data demonstrated that rotavirus infected mice had decreased diarrhea at 3 and 5 DPI, we utilized Balb/C mouse pup derived jejunal organoids to assess if inhibiting P2Y1 signaling in epithelial cells could suppress rotavirus induced fluid secretion. To test this, we used the organoid swelling assay^13, 21^, in which the increase in organoid cross-sectional area is measured to estimate the relative volume of fluid secreted into the organoid lumen (**Figure 3A**). In mock-infected organoids, MRS2500 treatment did not affect organoid swelling (**Figure 3B**). D6/2-infection resulted in a larger increase in the organoid cross-sectional area relative to mock-infected controls, while infected organoids treated with MRS2500 had a significantly dampened area increase (**Figure 3B**). These data suggest that the rotavirus-induced increase in P2Y1 signaling is involved in the increased epithelial fluid secretion and can contribute to rotavirus diarrhea severity.

**Figure 3:**
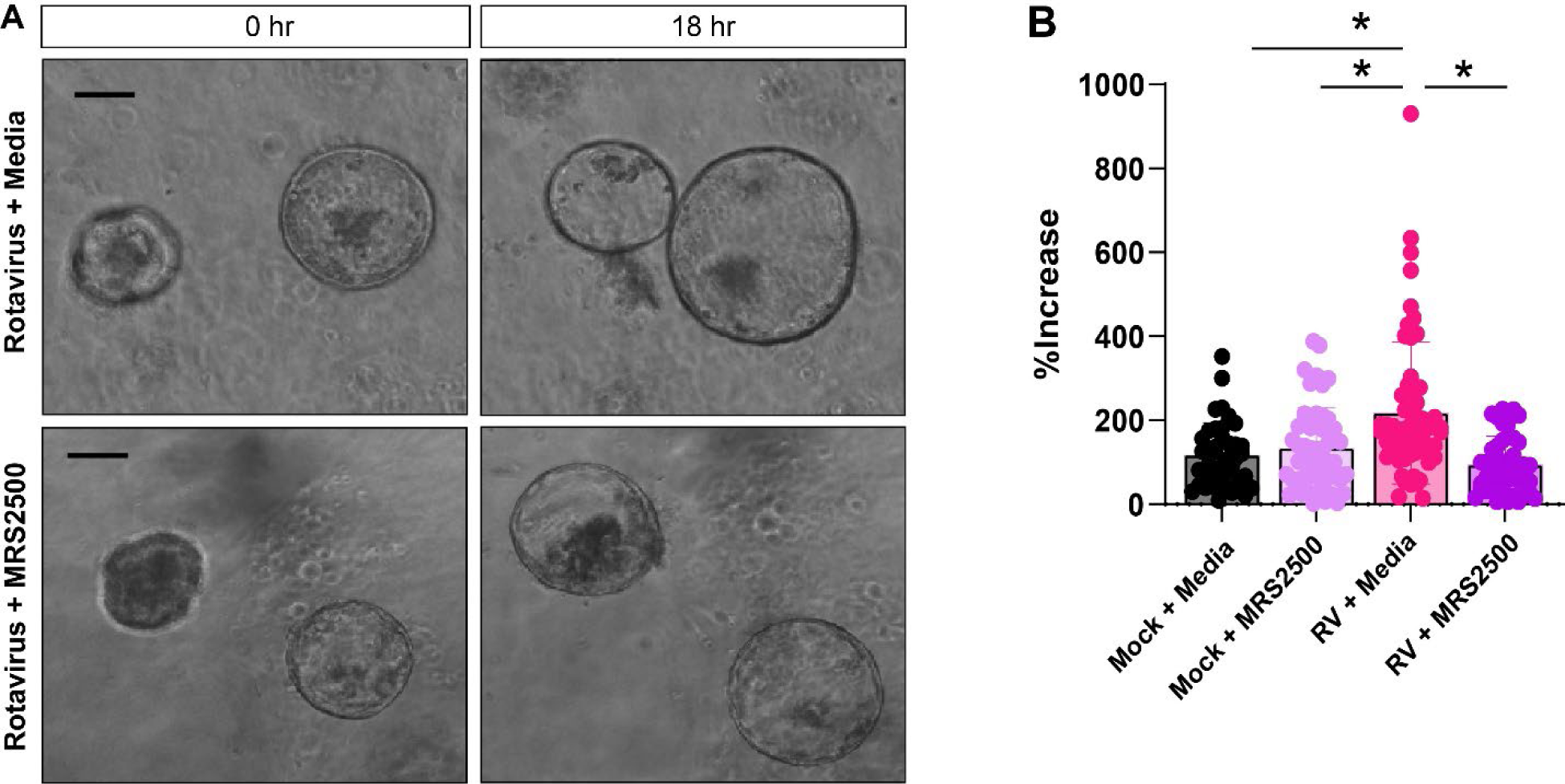
P2Y1 inhibition results in decreased fluid secretion in vitro. **A.** Representative images of organoid swelling assay at 0hr and 18hr post infection. 3D mouse jejunal organoids infected with D6/2 rotavirus and treated with PBS or MRS2500. Scale bar=100 μm. **B.** Percent increase between basal and maximum organoid swelling comparing mock uninfected (black) vs D6/2 infection (magenta) mouse organoids treated with PBS vs MRS2500 (purple). One Way ANOVA. *p<0.05.

In cell culture, we’ve demonstrated that P2Y1 signaling increased the rate of rotavirus spread^36^. As P2Y1 was able to minimize fluid secretion *in vitro* and decrease diarrhea severity, duration, and prevalence in mouse pups, we sought to identify the role of P2Y1 in regulating the levels of rotavirus in the intestine and shedding into the stool (**Figure 4**). Using immunofluorescence, we detected abundant rotavirus antigen at the tips of the intestinal villi at 1 DPI and up to 5DPI (**Figure 4A**). Quantification of our immunostaining revealed that there were fewer rotavirus infected cells in MRS2500 treated mice at 3 DPI compared to PBS-treated pups (**Figure 4B**). At 3 DPI, in PBS-treated pups, rotavirus-infected cells were observed in several villi, but in MRS2500-treated pups, there were fewer rotavirus positive cells and most villi were rotavirus negative. At 5 DPI, rotavirus was restricted to only a few villi in the PBS-treated pups and infrequently detected in the MRS2500-treated pups (**Figure 4A**). We also assessed rotavirus genome levels in the stool by qRT-PCR (**Figure 4C**). As expected, we found high levels of rotavirus at 1, 2, and 3 DPI in stool from rotavirus infected mice, with levels decreasing on 4 and 5 DPI. MRS2500 treated mice had lower levels of rotavirus in the stool at 4 and 5 DPI compared to saline treated mice, which represents levels after 3-4 treatments respectively (**Figure 4C**). These findings indicate that P2Y1 signaling contributes to rotavirus infection and shedding, and that blocking P2Y1 may affect the rate by which rotavirus-infected cells are cleared from the gastrointestinal tract.

**Figure 4:**
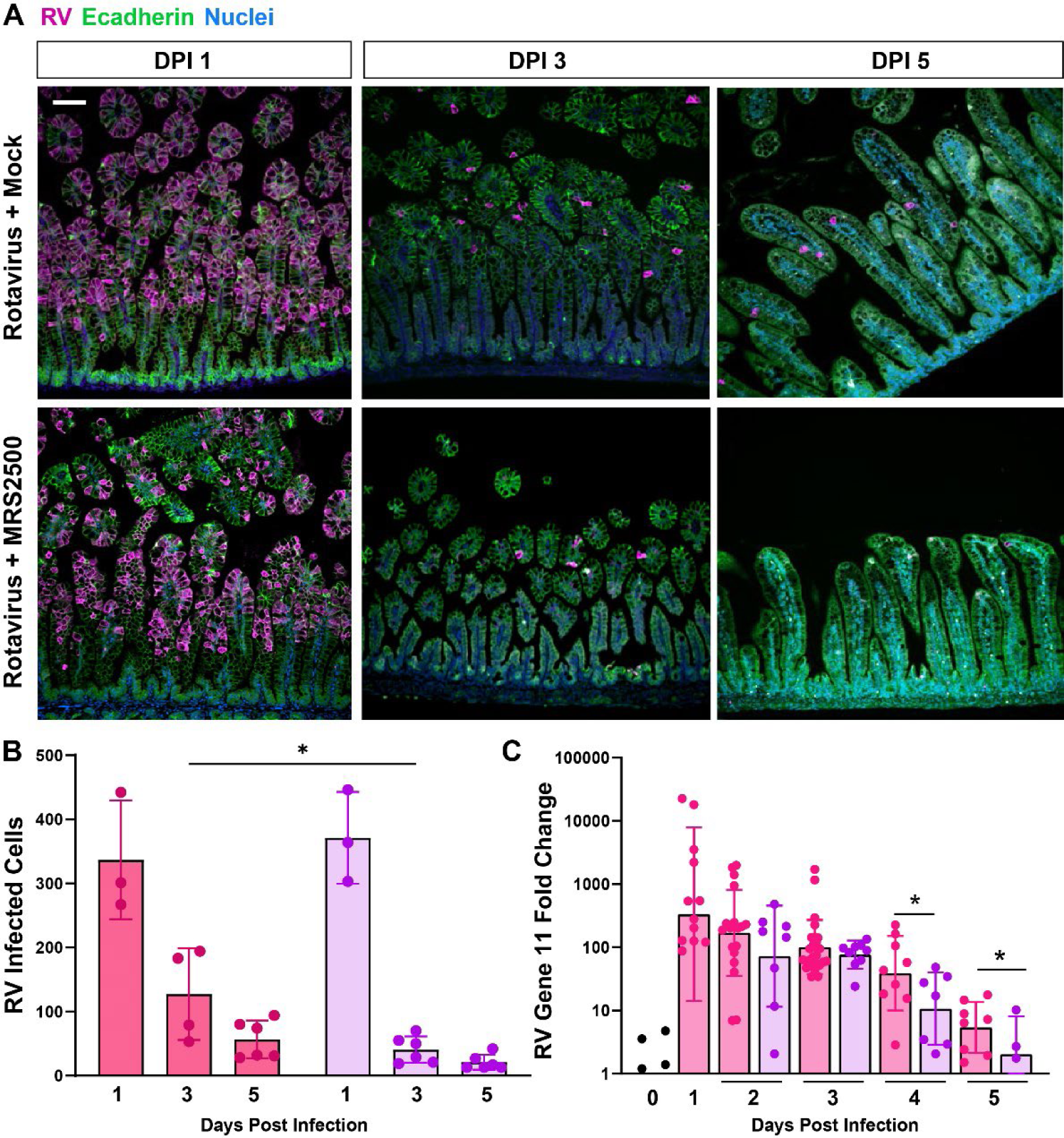
P2Y1 inhibition affects rotavirus replication and shedding. **A.** Representative images of D6/2 rotavirus immunostaining at 1, 3 and 5 DPI in the terminal ileum of rotavirus infected mice treated with PBS vehicle or MRS2500. Rotavirus is depicted in pink, E-cadherin is depicted in green, and nuclei are depicted in blue. Scale bar = 100 µm. **B.** Quantification of the total number of rotavirus infected cells in the intestine of D6/2 infected mice treated with PBS (magenta, n=3-6 pups) or MRS2500 (purple, n=3-6 pups) at 1, 3 and 5 DPI. Two Way ANOVA, *p <0.05. **C.** qPCR of stool from D6/2 infected pups treated with PBS (magenta, n= 6-18 stooled pups) or MRS2500 (purple, n=5-15 stooled pups) at 0-5 DPI for rotavirus gene 11 mRNA transcripts, normalized to 18S mRNA transcripts and fold change relative to 0 DPI. Multi-Repeated Measures ANOVA, *p <0.05. n=3-10 pups per treatment group.

Rotavirus is primarily transmitted through the fecal-oral route, and strategies to reduce transmission, particularly in the setting of an outbreak, are important for disease control. Based on our above findings that P2Y1 limits rotavirus diarrhea, as well as replication and virus shedding, we sought to determine whether inhibition of P2Y1 signaling in uninfected hosts could reduce rotavirus diarrhea during transmission. Murine rotavirus is efficiently transmitted from infected pups to uninfected cage-mates^37^, so we utilized this to examine if P2Y1 inhibition would alleviate symptoms of rotavirus transmission (ie diarrhea). For these studies, we used CD-1 mice because they are more susceptible to rotavirus than B57Bl6 mice and are outbred, better mimicking rotavirus transmission in a susceptible population. For each CD-1 mouse litter, 30% of pups were inoculated by oral gavage with D6/2 rotavirus, and the remaining 70% littermates were given daily oral gavage of 4mg/kg MRS2500 or sterile saline (PBS) starting at 0 DPI (**Figure 5A**). Similar to the studies above, diarrhea onset in rotavirus-infected pups was 2 DPI, with peak incidence from 3-5 DPI (**Figure 5B**). For the uninoculated littermates, diarrhea onset was delayed by 1 day for both the PBS- and MRS2500-treated pups and the overall prevalence and severity was less than that of the directly infected pups (**Figure 5B-C**). While uninoculated littermates in both treatment groups exhibited diarrhea (**Figure 5C**), treating the uninoculated pups with MRS2500 significantly reduced the diarrhea prevalence from ∼92% per litter to ∼56% per litter (**Figure 5D**). Rotavirus was detectable by fluorescent foci assay in stool regardless of diarrhea status or treatment group in uninoculated pups (**Supplemental Table 1**). While prophylactic inhibition of P2Y1 signaling prevents symptomatic rotavirus diarrhea in more than half of each litter, it does not appear to prevent rotavirus transmission. Collectively, these studies establish that rotavirus exploitation of P2Y1 signaling is a significant contributor to rotavirus diarrhea through increased virus-induced fluid secretion, virus replication and shedding.

**Figure 5:**
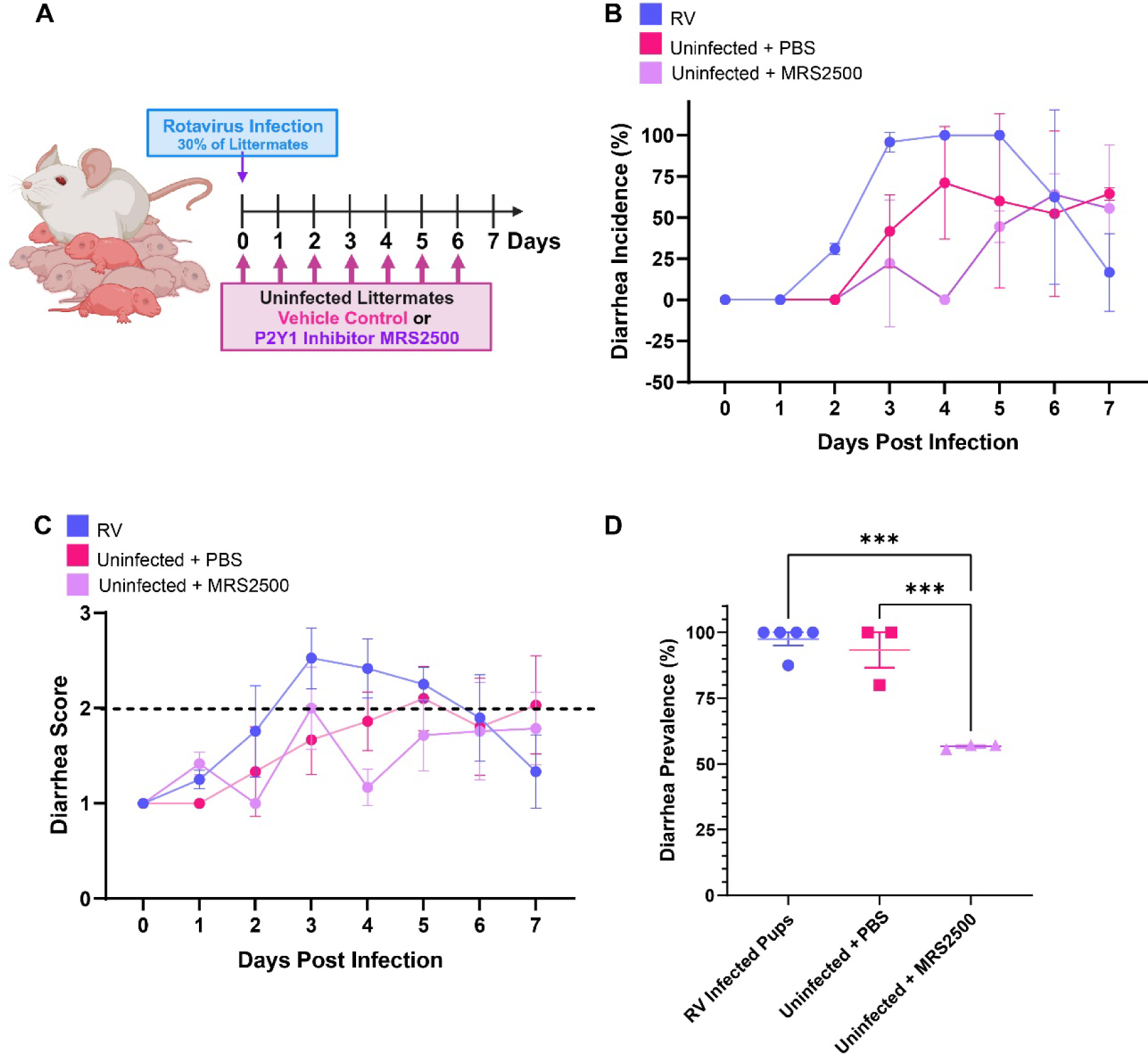
P2Y1 inhibitor treatment decreases effects of rotavirus transmission. **A.** Schematic diagram of our experimental setup. In CD-1 dams with natural litters, 30% of pups were orally gavaged with rotavirus at day 4-6 of life and monitored daily. Uninfected littermates were given either vehicle control (PBS) or 4mg/kg MRS2500 (P2Y1 inhibitor) by oral gavage daily, starting at 0DPI, until euthanasia. Stool and diarrhea scores were assessed and collected daily from infected and uninfected littermates. **B.** The average percentage of CD-1 pups with diarrhea as recorded every 24 hours for rotavirus infected pups (n=6 litters; blue), uninfected littermates treated with PBS (n=3 litters, magenta), or uninfected littermates treated with MRS2500 (n=3 litters, purple). **C.** Diarrhea scores over a 7-day period from rotavirus infected pups (n=6 litters; blue), uninfected littermates treated with PBS (n=3 litters, magenta), or uninfected littermates treated with MRS2500 (n=3 litters, purple). **D**. Diarrhea prevalence over 7-day period in rotavirus inoculated pups and uninfected pups treated with PBS or MRS2500. One-Way ANOVA. ***p<0.005.

## Discussion

We previously discovered that rotavirus infected cells release ADP, which initiates large scale intercellular calcium waves through activation of P2Y1 purinergic receptors on neighboring cells, making this an important contributor to cell-cell communication during infection^21^. We also recently performed a comprehensive survey of the purinergic receptor expression by gastrointestinal epithelial cells and found that of the 15 purinergic receptors, P2Y1 shows the most functional activity throughout the gastrointestinal tract^38^, indicating that exploitation of P2Y1 signaling by rotavirus could have broad-ranging repercussions to rotavirus infection and disease severity. Using the neonatal mouse model of rotavirus diarrhea, we examined the effect of blocking P2Y1 signaling on virus replication and diarrhea and found that P2Y1 signaling influences multiple aspects of rotavirus disease, including diarrhea prevalence, fluid secretion, and virus clearance. Together these data demonstrate that P2Y1 signaling is integral for rotavirus infection and disease, establishing P2Y1 as a potential target for anti-diarrheal therapies.

Rotavirus diarrhea is multifactorial; as a non-inflammatory diarrhea, it has been attributed to several mechanisms including osmotic diarrhea due to malabsorption (by enterocyte destruction)^39–41^ and secretory diarrhea due to enteric nervous system activation^7, 24, 42^. Increased chloride secretion is a significant contributor to rotavirus-induced fluid loss, so identifying the pathways that increase chloride secretion during rotavirus infection has long been considered an attractive target for new anti-diarrheal drugs. This study shows that P2Y1 activation plays an important role in rotavirus diarrhea severity *in vivo* and this correlates with increased rotavirus-induced fluid secretion *in vitro*. This finding aligns well with single cell RNA sequencing data which indicates small intestinal epithelial cells, including chloride (Cl-) secreting enterocytes, express P2Y1^43^. Purinergic receptors have been established to play an important role in regulating Cl^-^ secretion in different epithelial cell types, including colonic epithelial cells^44–46^. It is interesting to note that in Ussing chamber studies of both Caco-2 and T84 colonic adenocarcinoma cell lines, the P2Y1 agonist ADP was a relatively weak activator of Cl^-^ secretion^46^. Yet, this is consistent with our recent studies showing that these cancer cell lines exhibit significantly lower functional P2Y1 expression than primary intestinal epithelial cells^38^. Additionally, in murine organoids, blocking P2Y1 inhibited rotavirus-induced organoid swelling, which is a measure of fluid secretion, but did not affect fluid secretion by mock-infected organoids. This suggests that P2Y1-induced fluid secretion was not a major pathway for basal fluid secretion but, as indicated by our imaging studies, may be selectively upregulated in response to infection.

The above data raises the question of what Cl^-^ secretion pathway is activated by rotavirus induced P2Y1 signaling. Most of the purinergic receptors expressed by gastrointestinal epithelial cells are coupled to Gq-containing heterotrimeric G-proteins and as such, activate phospholipase C (PLC) and inositol trisphosphate (IP3)-dependent mobilization of ER Ca^2+^, which implicates Cl^-^ secretion by Ca^2+^-activated Cl^-^ channels (CaCCs). This is supported by studies showing that purinergic receptor mediated Cl^-^ secretion is blocked by pretreatment with phospholipase C (PLC) inhibitors or Ca^2+^ chelators, indicating elevation of cytosolic Ca^2+^ is critical for the downstream activation of Ca^2+^ activated Cl^-^ channels (CaCCs), such as anoctamin 1 (Ano1, aka TMEM16A)^44^. Studies have implicated Ano1 as a major source of epithelial Cl^-^ secretion during rotavirus induced diarrhea; however, the functional level of Ano1 in the small intestine remains controversial and there is evidence to suggest Ano1 is not expressed in normal intestinal epithelial cells^32, 33,34, 35, 47–49^. Thus, future work is needed to identify the source(s) of Cl-secretion that contributes to rotavirus diarrhea. Nevertheless, our results demonstrate that P2Y1 signaling is a key part of rotavirus-induced fluid secretion and determining the Cl-channel(s) activated downstream of P2Y1 will help unravel pathways involved in the secretory component of rotavirus diarrhea.

A separate, but related, contributor to rotavirus-induced fluid secretion is serotonin release from enterochromaffin cells^50, 51^. Previous work has shown that serotonin receptor antagonists or broad inhibition of the enteric nervous system lessen the severity of rotavirus-associated disease in mice^42, 51^. However, directly targeting serotonin release in children may have detrimental effects on development. We previously reported that P2Y1 signaling promotes human rotavirus-induced serotonin release in human intestinal organoids^21^. Previous studies have found that purine release upon mechanostimulation of epithelial cells can trigger serotonin secretion from enterochromaffin cells through activation of P2Y1^52^. Similarly, our model is that ADP released from rotavirus-infected enterocytes activates P2Y1 on enterochromaffin cells in a paracrine manner, resulting in an increase in cytosolic Ca^2+^ that triggers serotonin release^21^. Thus, inhibition of serotonin secretion by blocking P2Y1 may have contributed to the observed decrease in diarrhea in pups treated with MRS2500. Given the prominent role of P2Y1 in GI epithelial purinergic signaling, dissecting the relative contribution of individual pathways (e.g., direct activation of CaCCs versus serotonin-mediated activation of the ENS) remains a complex challenge. Future studies using epithelial cell-type-specific conditional knockouts of P2Y1 expression will be needed to assess the importance of distinct P2Y1-activated pathways in the overall pathogenesis of rotavirus.

Beyond the intestinal epithelium, small molecule P2Y1 blockers, like MRS2500, could reduce rotavirus diarrhea by directly inhibiting P2Y1 signaling in enteric neurons^53^ and/or smooth muscle^54^. Other studies have shown that P2Y1 activation in colonic smooth muscle contributes to hyperpolarization and relaxation^55, 56^ yet it remains unclear the contribution purinergic signaling has as a neurotransmitter to affect smooth muscle relaxation. While further work would be needed to assess the broader effects of P2Y1 inhibition in our model, it is important to note that we did not observe changes in stooling in MRS2500 treatment groups of infected or uninfected pups. Thus, in the absence of rotavirus-induced diarrhea, MRS2500 did not markedly affect stool frequency.

One interesting observation from this study was that inhibiting P2Y1 limited rotavirus infection in the villi and reduced rotavirus shedding in the feces starting around 3DPI. We have previously shown that rotavirus is capable of infecting P2Y1 deficient MA104 cells and human intestinal organoids to the same degree as parental lines harboring P2Y1, demonstrating an equivalent susceptibility of these cells to rotavirus infection^21^. We have also demonstrated that rotavirus is able to infect LLC-MK2 cells which do not express P2Y1; however, over-expressing P2Y1 in LLC-MK2 cells enhanced rotavirus spread^36^. Thus, we do not anticipate that the lower levels of rotavirus observed in mice treated with MRS2500 at 3 and 5 DPI reflect insusceptibility to rotavirus. The decreased presence of rotavirus at 3DPI in MRS2500 treated pups may be interconnected with the observed decreased diarrhea severity and incidence beginning at 3DPI. It is possible that the decreased rotavirus infection in MRS2500 treated pups leads to the decreased diarrhea observed, not only because of P2Y1 signaling inhibition but also due to fewer infected cells. However, our organoid swelling data suggests that decreased rotavirus spread is not the only means by which P2Y1 inhibition limits symptoms of rotavirus infection.

Our data demonstrates that prophylactic MRS2500 treatment of uninfected littermates significantly decreases diarrhea prevalence but does not fully prevent diarrhea from occurring. However, prophylactic MRS2500 treatment in uninfected littermates did not prevent rotavirus infection as we were able to detect rotavirus in their stool. This suggests that the effects of P2Y1 inhibition are primarily through limiting epithelial (and epithelial-triggered ENS)-mediated responses to rotavirus infection, which includes chloride secretion and serotonin release, that contribute to secretory diarrhea. As blocking P2Y1 reduced rotavirus spread, we expect this to contribute to the decreased prevalence of diarrhea by limiting multi-round virus replication and the resulting amplification of epithelial dysregulation.

Ultimately, we found that blocking P2Y1 signaling *in vivo* significantly limits rotavirus infection and diarrhea, which shows that P2Y1 may be an effective target for development of a dual function anti-diarrheal and antiviral therapy for short-term treatment of an acute rotavirus infection. While we did not completely prevent rotavirus diarrhea in these studies, we achieved a ∼25% reduction in diarrhea prevalence with MRS2500 treatment started on 1-day post-infection. There are several potential explanations for this observation. One possibility is the involvement of other P2Y1-independent pathways or mechanisms which contribute to rotavirus diarrhea. Another possibility is that increased P2Y1 signaling during the first day of infection induces strong enough epithelial dysfunction, including fluid secretion and activation of the ENS, such that even upon blocking P2Y1, diarrhea continues for some time. Alternatively, the pharmacokinetics of MRS2500 in the intestine, particularly in pups, is not well known, and even with daily gavage, inhibition of P2Y1 signaling may wane between treatments. Thus, further studies with different doses and treatment schedules will refine the optimal therapeutic approach to eliminating rotavirus diarrhea by blocking P2Y1. In conclusion, our data highlight the importance of P2Y1 in rotavirus infection and suggest that P2Y1 could serve as a novel target for treating rotavirus infection.

---

**Supplemental Table 1:** Detection of Rotavirus in Stool (by FFA) of Diarrheal Scored CD-1 Pups Pretreated with PBS or MRS2500 During Rotavirus Transmission Studies.

|  | No Diarrhea |  | Diarrhea |  |
| --- | --- | --- | --- | --- |
|  | Not Detected | RV Detected | Not Detected | RV Detected |
| **PBS** | 16.4% ± 11.5 | 23.9% ± 19.4 | 31.9% ± 21.8 | 48.7 ± 21.7 |
| **MRS2500** | 37.9% ± 14.1 | 27.8% ± 16 | 7.14% ± 0 | 48.1% ± 11.9 |

